## Supplemental Figures for "Repurposing of KLF5 activates a cell cycle signature during the progression from a precursor state to Oesophageal Adenocarcinoma"

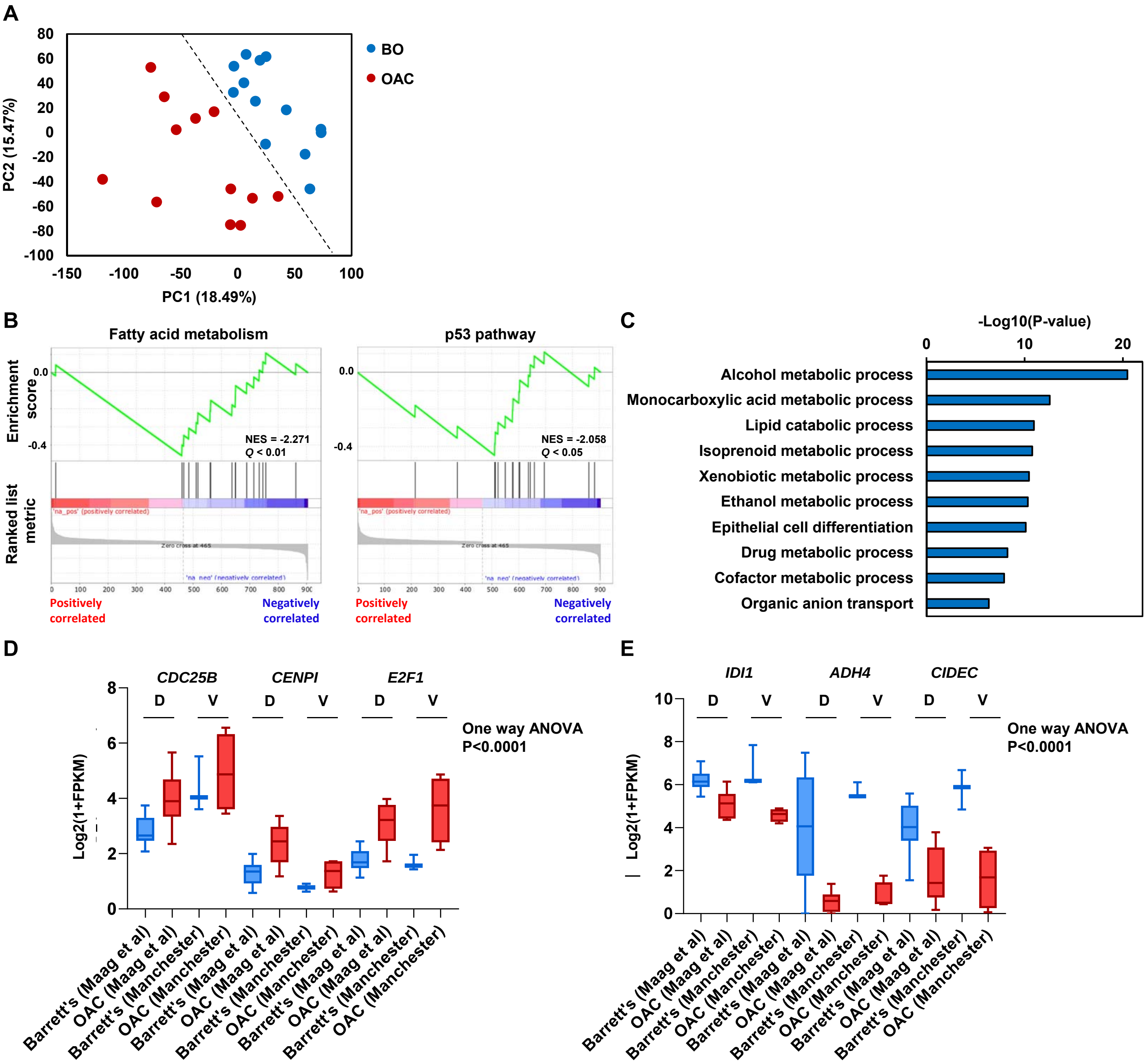


**Figure 1- supplement 1. Differential gene expression analysis between Barrett's and OAC patient samples.** (A) PCA plot of 13 Barrett's oesophagus (blue) and 12 OAC (red) RNA-seq samples (Maag et al., 2017). (B) GSEA plots of significantly downregulated gene sets in OAC compared to Barrett's oesophagus. Normalised enrichment score (NES) and *Q*-value are shown. (C) Biological pathway GO term of significantly downregulated genes in OAC compared to Barrett's oesophagus. (D) Box plot of $Log_2$(1+FPKM) expression from BO (blue) and OAC (red) samples in discovery (D; Maag et al., 2017) and validation (V; this study) datasets for upregulated example genes (D) and downregulated example genes (E). Whiskers represent minimum and maximum values and one way ANOVA *p*-value is shown.

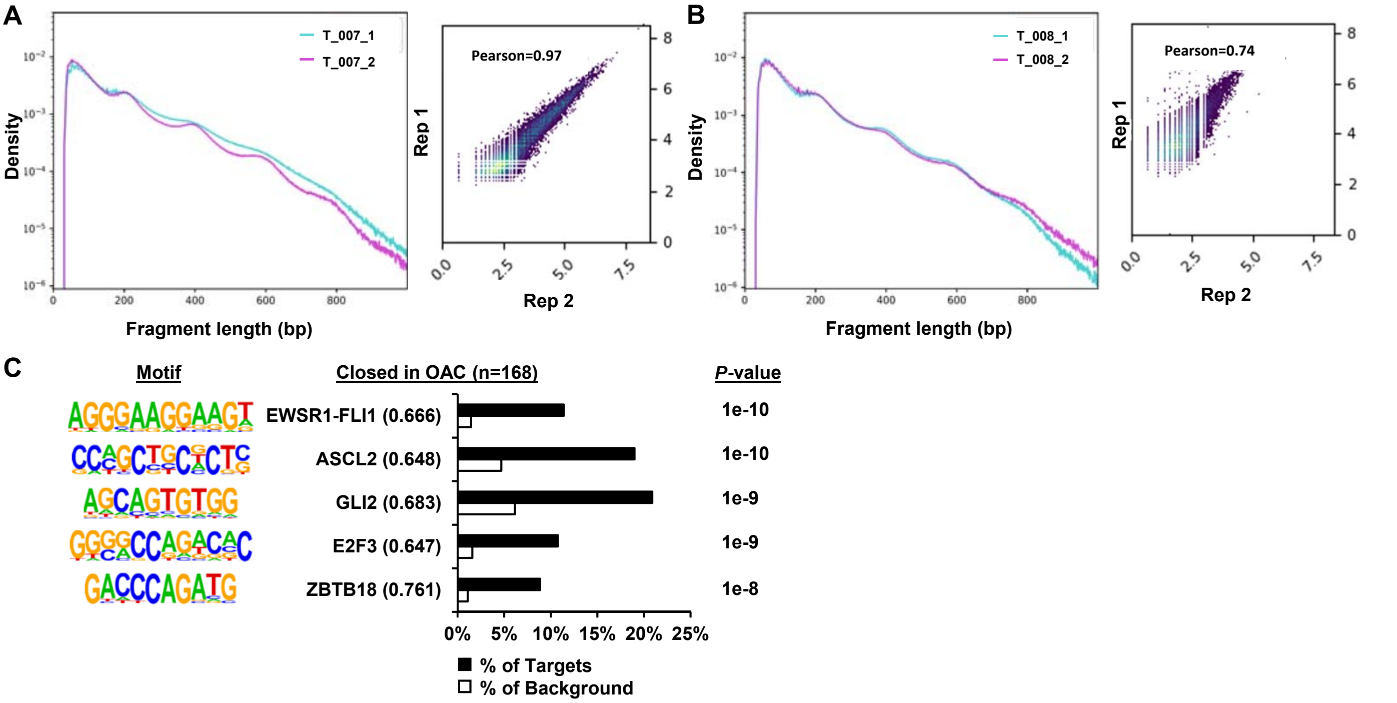


**Figure 2- figure supplement 1. ATAC-seq analysis of patient OAC samples.** (A) Fragment length plot of $log_{10}$ transformed ATAC-seq signal from technical replicates 1 and 2 of the T_007 OAC sample (left). Correlation plot of ATAC-seq signal from technical replicates 1 and 2 of the T_007 OAC sample at ATAC-seq peaks called from replicate 1 (right). Pearson correlation coefficient is shown. (B) Fragment length plot of $log_{10}$ transformed ATAC-seq signal from technical replicates 1 and 2 of the T_008 OAC sample (left). Correlation plot of ATAC-seq signal from technical replicates 1 and 2 of the T_008 OAC sample at ATAC-seq peaks called from replicate 1 (right). Pearson correlation coefficient is shown. (C) Bar chart of percentage targets and percentage background of *de novo* discovered motifs at regions with significant decreased chromatin accessibility in OAC. *De novo* motifs, called transcription factor with motif match score (in brackets) and *P*-value are shown.

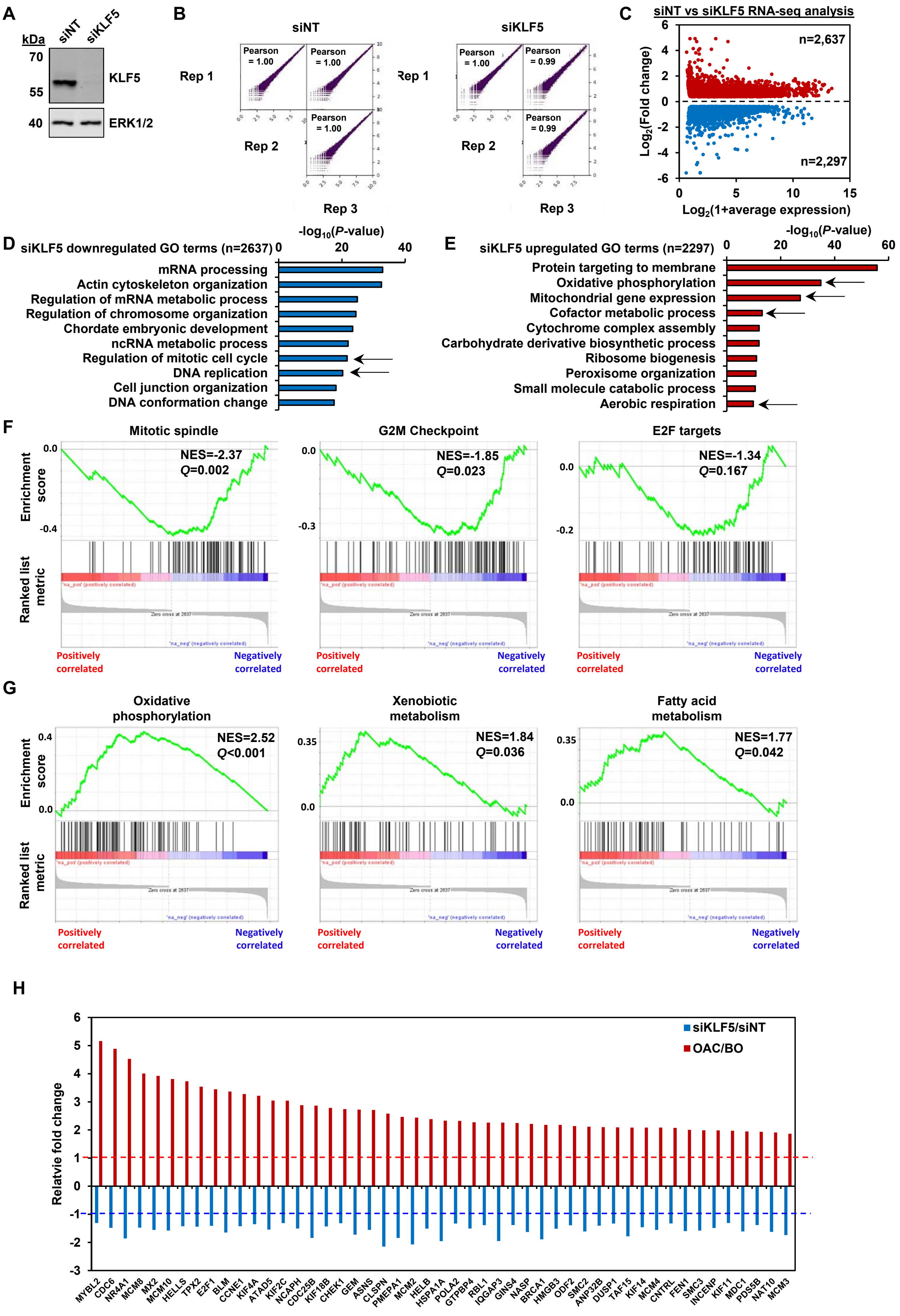

A
kDa
siNT
siKLF5
70
55
KLF5
40
ERK1/2
B
siNT
siKLF5
Rep 1
Rep 2
Rep 3
Pearson = 1.00
Pearson = 0.99
C
siNT vs siKLF5 RNA-seq analysis
n=2,637
n=2,297
Log2(Fold change)
Log2(1+average expression)
D
siKLF5 downregulated GO terms (n=2637)
-log10(P-value)
mRNA processing
Actin cytoskeleton organization
Regulation of mRNA metabolic process
Regulation of chromosome organization
Chordate embryonic development
ncRNA metabolic process
Regulation of mitotic cell cycle
DNA replication
Cell junction organization
DNA conformation change
E
siKLF5 upregulated GO terms (n=2297)
-log10(P-value)
Protein targeting to membrane
Oxidative phosphorylation
Mitochondrial gene expression
Cofactor metabolic process
Cytochrome complex assembly
Carbohydrate derivative biosynthetic process
Ribosome biogenesis
Peroxisome organization
Small molecule catabolic process
Aerobic respiration
F
Mitotic spindle
NES=-2.37
Q=0.002
G2M Checkpoint
NES=-1.85
Q=0.023
E2F targets
NES=-1.34
Q=0.167
Enrichment score
Ranked list metric
Positively correlated
Negatively correlated
G
Oxidative phosphorylation
NES=2.52
Q<0.001
Xenobiotic metabolism
NES=1.84
Q=0.036
Fatty acid metabolism
NES=1.77
Q=0.042
H
siKLF5/siNT
OAC/BO
Relatvie fold change
MYBL2 CDC6 NR4A1 MCM8 MX2 MCM10 HELLS TPX2 E2F1 BLM CCNE1 KIF4A ATAD5 KIF2C NCAPH CDC25B KIF18B CHEK1 GEM ASNS CLSPN PMEPA1 MCM2 HELB HSPA1A POLA2 GTPBP4 RBL1 IQGAP3 GINS4 NASP BRCA1 HMGB3 ODF2 SMC2 ANP32B DUSP1 TAF15 KIF14 MCM4 CNTRL FEN1 SMC3 INCENP KIF11 MDC1 PDS5B NAT10 MCM3

**Figure 3- figure supplement 1. Identification of the KLF5-regulated cistrome in OE19 cells.** (A) Immunoblot of protein lysate from OE19 cells treated with either siNT or siKLF5 and probed with antibodies against KLF5 and ERK1/2. (B) Correlation plots of RNA-seq data from three biological replicates. Pearson correlations are shown. (C) Scatter plot of significantly differentially expressed genes (red are up- and blue are down-regulated genes; ±1.3x, *Q*-value<0.05) in OE19 cells with siKLF5 treatment. (D) Biological processes GO term analysis of downregulated genes with siKLF5 treatment. Cell cycle related GO terms are indicated with arrows. (E) Biological processes of GO term analysis of upregulated genes with siKLF5. Metabolism related GO terms are indicated with arrows. (F) Gene set enrichment analysis of downregulated genes with siKLF5 treatment. Cell cycle related gene sets, normalised enrichment scores (NES) and *Q*-values are shown. (G) Gene set enrichment analysis of upregulated genes with siKLF5 treatment. Metabolism related gene sets, normalised enrichment scores (NES) and *Q*-values are shown. (H) Bar chart of fold change in expression of cell cycle related genes in siKLF5 treated OE19 cells (blue; n=3) or patient OAC samples (red

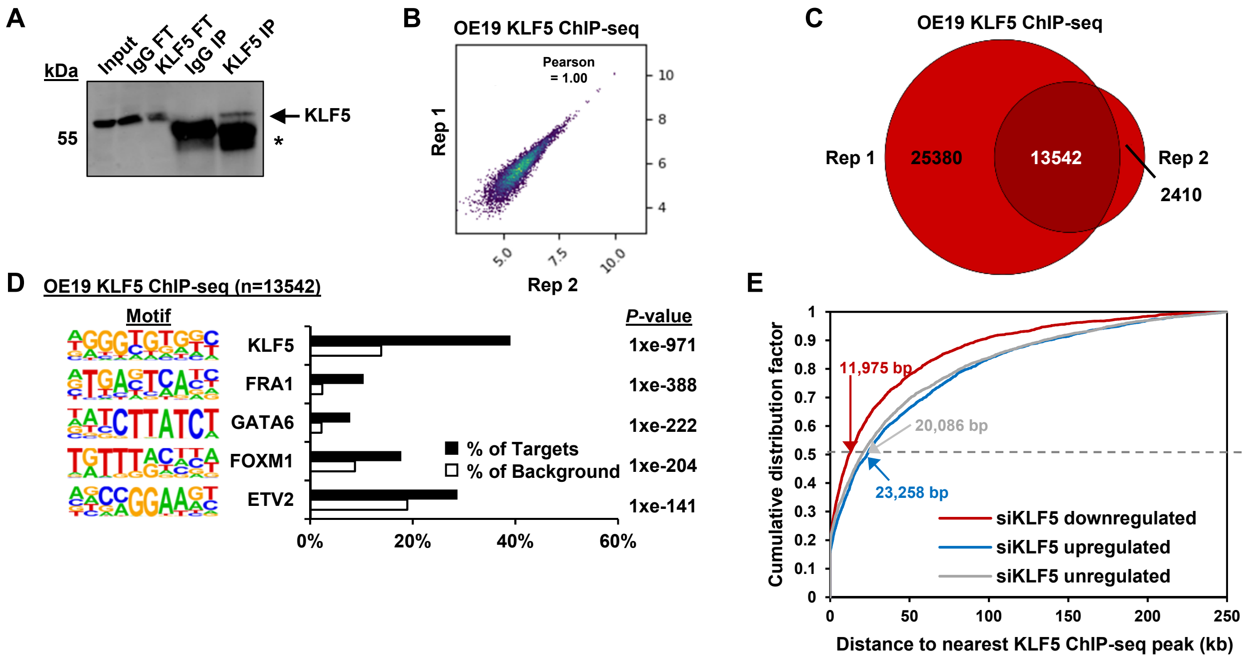


**Figure 3- figure supplement 2. KLF5 ChIP-seq analysis in OE19 cells.** (A) Immunoblot of KLF5 from OE19 cells following ChIP in different fractions; IgG FT (flow through from non-specific IgG precipitation), KLF5 FT (flowthrough from KLF5 precipitation), IgG elute (material eluted following IgG precipitation) KLF5 IP (material eluted following KLF5 immunoprecipitation). Arrows indicating immunoprecipitated protein (KLF5) and asterisks are IgG heavy chain. (B) Correlation plot of KLF5 ChIP-seq data from OE19 cells from two biological replicates. Pearson correlation coefficient is shown. (C) Venn diagrams showing overlap of peaks called by each replicate (Rep) for KLF5 ChIP-seq from OE19 cells. The numbers of peaks in each sector are given. (D) Bar chart of percentage targets and percentage background of *de novo* discovered motifs in OE19 KLF5 ChIP-seq peaks. *De novo* motif, called transcription factor with motif match score (in brackets) and *P*-value are shown. Note that the KLF5 motif is shown in the reverse orientation to those in other figures). (E) Cumulative distribution of the closest KLF5 ChIP-seq peaks to the TSS of genes that were significantly down or upregulated following siKLF5 treatment compared to expressed genes (FPKM>1) that were not significantly deregulated by siKLF5 treatment. Median distances to the TSS are shown.

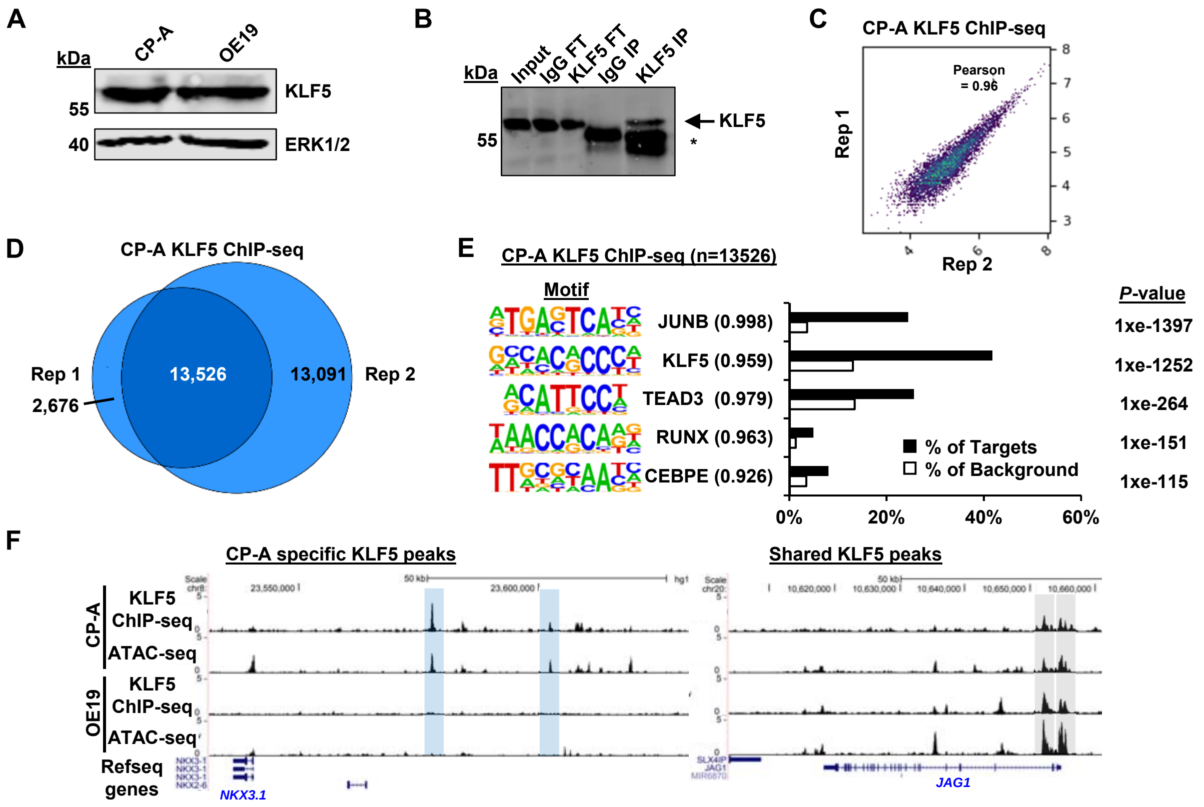


**Figure 4-figure supplement-1. KLF5 ChIP-seq analysis in CP-A cells.** (A) Immunoblot of CP-A and OE19 protein lysate probed with antibodies against KLF5 and ERK1/2. (B) Immunoblot of KLF5 from CP-A cells following ChIP in different fractions; IgG FT (flow through from non-specific IgG precipitation), KLF5 FT (flowthrough from KLF5 precipitation), IgG elute (material eluted following IgG precipitation) KLF5 IP (material eluted following KLF5 immunoprecipitation). Arrows indicating immunoprecipitated protein (KLF5) and asterisks are IgG heavy chain. (C) Scatter plots of KLF5 ChIP-seq data from CP-A from two biological replicates. Pearson correlation coefficient shown. (D) Venn diagrams showing overlap of peaks called by each replicate for KLF5 ChIP-seq from CP-A cells. The numbers of peaks in each sector are given. (E) Bar chart of percentage targets and percentage background of *de novo* discovered motifs at CP-A KLF5 ChIP-seq peaks. *De novo* motif, called transcription factor with motif match score (in brackets) and *P*-value shown. (F) Example UCSC genome browser tracks of KLF5 ChIP-seq and ATAC-seq data from CP-A and OE19 cells at the *NKX3-1* (left) and *JAG1* loci (right). CP-A specific peaks are highlighted in blue and shared peaks are highlighted in grey.

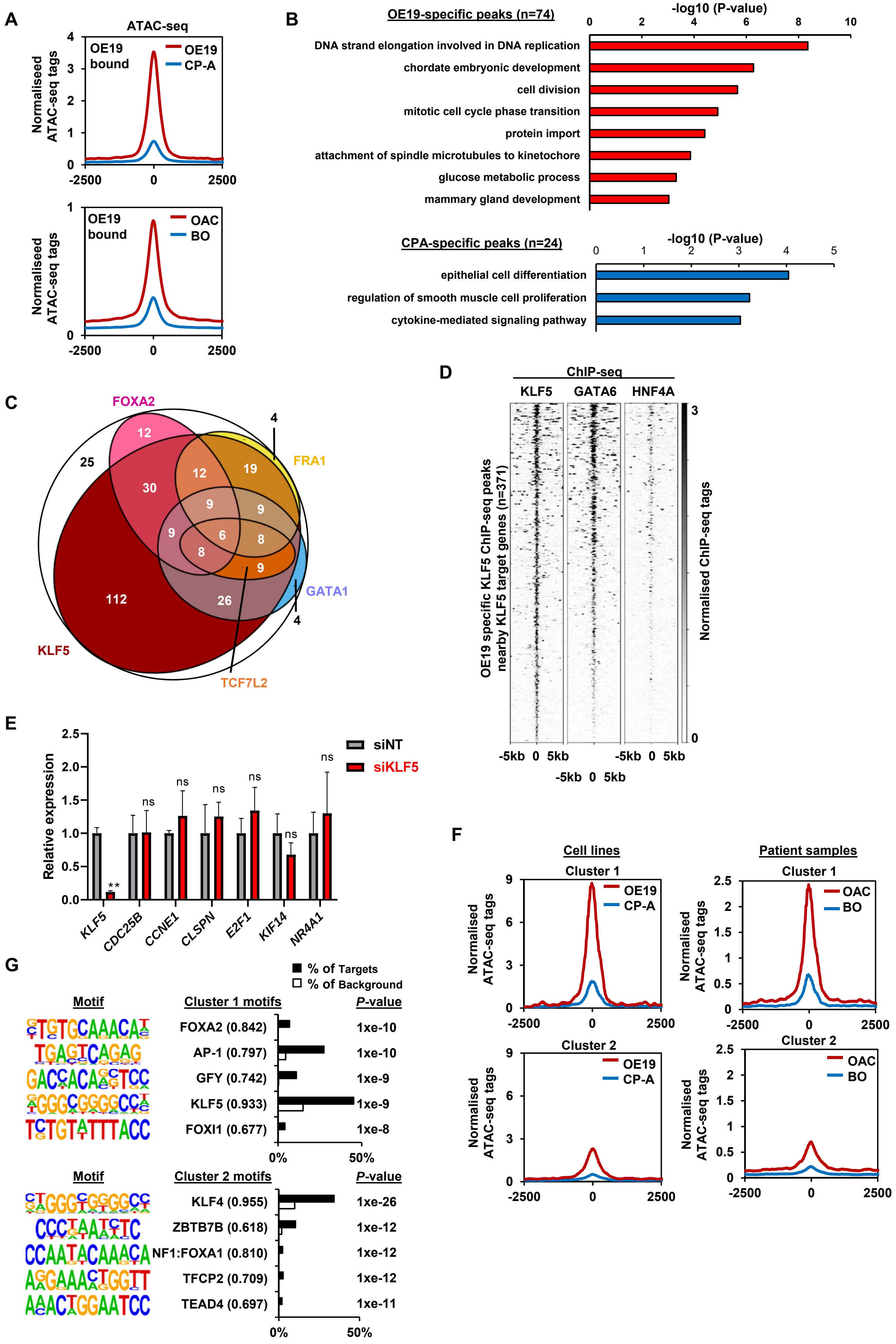

A
ATAC-seq
Normaliseed ATAC-seq tags
OE19 bound
OE19
CP-A
-2500
0
2500
OE19 bound
OAC
BO
B
OE19-specific peaks (n=74)
-log10 (P-value)
DNA strand elongation involved in DNA replication
chordate embryonic development
cell division
mitotic cell cycle phase transition
protein import
attachment of spindle microtubules to kinetochore
glucose metabolic process
mammary gland development
CPA-specific peaks (n=24)
-log10 (P-value)
epithelial cell differentiation
regulation of smooth muscle cell proliferation
cytokine-mediated signaling pathway
C
FOXA2
FRA1
GATA1
KLF5
TCF7L2
D
ChIP-seq
KLF5
GATA6
HNF4A
OE19 specific KLF5 ChIP-seq peaks nearby KLF5 target genes (n=371)
Normalised ChIP-seq tags
-5kb 0 5kb
E
Relative expression
siNT
siKLF5
KLF5
CDC25B
CCNE1
CLSPN
E2F1
KIF14
NR4A1
ns
**
F
Cell lines
Patient samples
Cluster 1
Cluster 2
Normalised ATAC-seq tags
OE19
CP-A
OAC
BO
G
% of Targets
% of Background
Motif
Cluster 1 motifs
P-value
FOXA2 (0.842)
1xe-10
AP-1 (0.797)
1xe-10
GFY (0.742)
1xe-9
KLF5 (0.933)
1xe-9
FOXI1 (0.677)
1xe-8
0%
50%
Cluster 2 motifs
KLF4 (0.955)
1xe-26
ZBTB7B (0.618)
1xe-12
NF1:FOXA1 (0.810)
1xe-12
TFCP2 (0.709)
1xe-12
TEAD4 (0.697)
1xe-11

**Figure 4-figure supplement-2. Integrative analysis of KLF5 ChIP-seq data in OE19 and CP-A cells.** (A) Tag density plots of normalised ATAC-seq signal at OE19 specific KLF5 ChIP-seq regions from CP-A (blue) and OE19 (red) cells (top) and merged BO (blue) and merged OAC (red) tissue (bottom). (B) Top GO terms of nearest genes associated with OE19-specific (top) or CP-A-specific (bottom) KLF5 peaks which show either increased (top) or decreased (bottom) expression in OAC. (C) Euler diagram of 299 KLF5 binding regions that are specific to OE19-specific binding regions that are located within loci (+/-250 kb) containing genes upregulated in OAC and downregulated with KLF5 depletion. The motifs identified in Fig. 4E (KLF5, GATA1, FOXA2, FRA1 and TCF7L2) found within each peak are shown (note that an additional 71 regions cannot be depicted due to the small numbers involved). The regions in the white circle contain none of the indicated motifs. (D) Heatmap of KLF5, GATA6 and HNF4A ChIP-seq signal from OE19 cells at OE19 specific KLF5 ChIP-seq peaks nearby KLF5 activated target genes. (E) RT-qPCR analysis of the indicated genes following treatment of CP-A cells with non-targeting (NT) or KLF5 siRNAs for 72 hours (n=3; P-values **=<0.01; ns=not significant). (F) Tag density plots of normalised ATAC-signal at cluster 1 (left) and cluster 2 (right) regions (see Fig. 4G) from CP-A (blue) and OE19 (red) cells (top) and merged BO (blue) and merged OAC (top) samples (bottom). (G) Bar chart of percentage targets and percentage background of *de novo* discovered motifs in cluster 1 (top) and cluster 2 (bottom) regions. *De novo* motif, called transcription factor with motif match score and *P*-value are shown

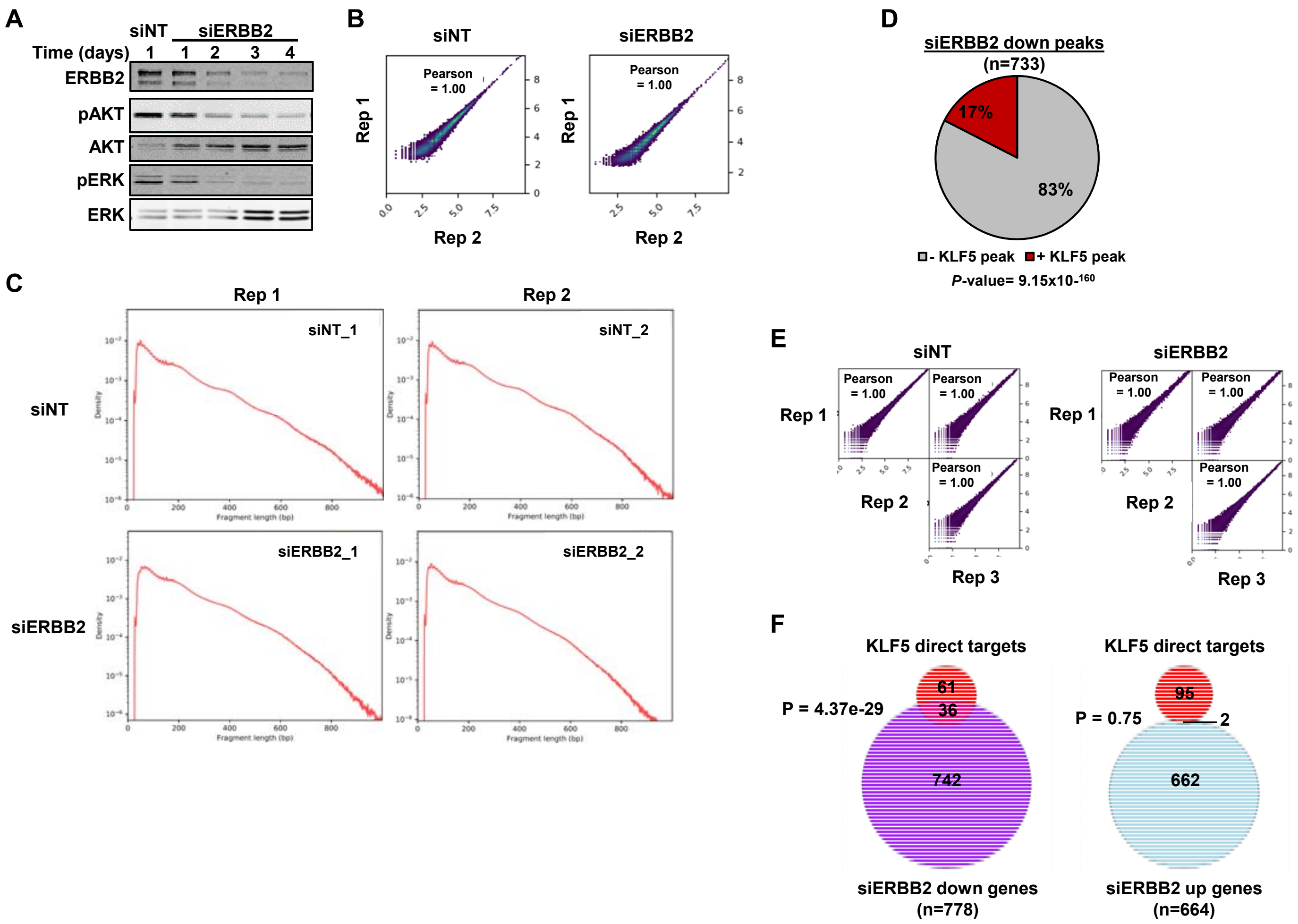


**Figure 5-figure supplement 1. ERBB2 and KLF5 regulate an overlapping set of genes.** (A) Immunoblot of OE19 protein lysate from cells treated with siERBB2 for up to 4 days or siNT probed with antibodies against ERBB2, phosphorylated AKT (pAKT), AKT, phosphorylated ERK (pERK) and ERK. (B) Correlation plots of replicates of ATAC-seq data from OE19 cells treated with either siNT or siERBB2 for 72 hours. Pearson correlations are shown. (C) Fragment length plots of replicates of ATAC-seq data from OE19 cells treated with either siNT or siERBB2. (D) Pie chart showing the percentage of ATAC-seq peaks showing a reduction of chromatin accessibility upon siERBB2 treatment that overlap with a KLF5 ChIP-seq peak in OE19 cells. Percentages and P-value (Fisher exact) for the overlap are shown. (E) Correlation plots of replicates of RNA-seq data from OE19 cells treated with either siNT and siERBB2 for 72 hours. Pearson correlations are shown. (F) Venn diagrams showing the overlap between the 97 direct KLF5 activated genes and genes either down- (left) or up- (right) regulated following siERBB2 treatment of OE19 cells (2 fold change, FDR<0.05, FPKM>1). P-values are hypergeometric tests.

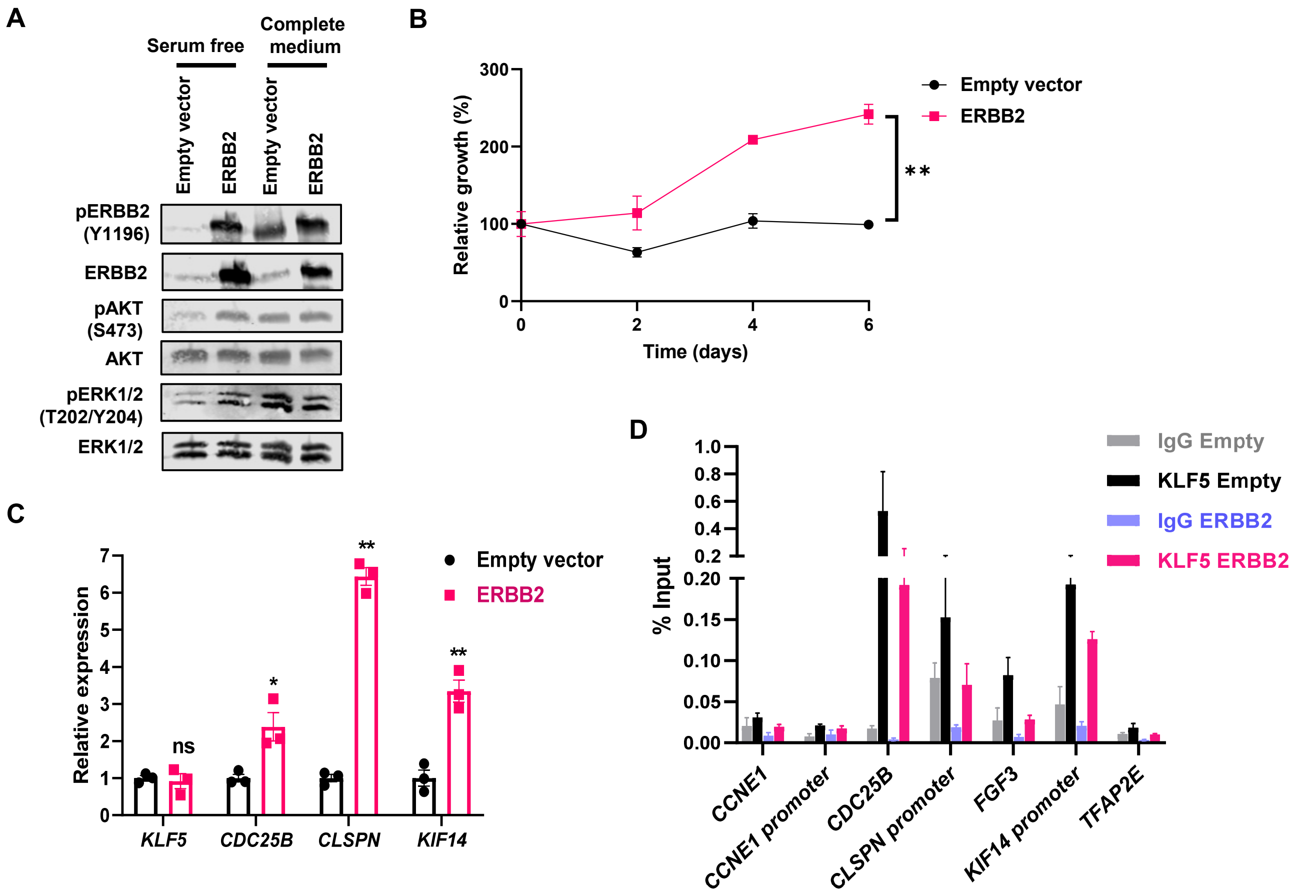


**Figure 5-figure supplement 2. ERBB2 overexpression drives growth factor independent proliferation and gene expression in BO-derived CP-A cells.** (A) Western blots of the indicated proteins or phosphorylated (p) sites in CP-A cells stably transfected with empty or ERBB2 expressing vectors grown in serum free medium for 48 hrs (lanes 1 and 2) or serum free media and stimulated with complete media for 15 mins (lanes 3 and 4). (B) Relative growth of CP-A cells stably transfected with empty or ERBB2 expressing vectors grown in serum free media for 6 days (n=3; **= P-value <0.01, two way ANOVA). (C) RT-qPCR analysis of the indicated genes (normalised to *GAPDH*) in serum starved CP-A cells stably transfected with empty or ERBB2 expressing vectors (n=3; P-values, *=<0.05, **= <0.01). (D) ChIP-qPCR of KLF5 binding to regulatory regions associated with the indicated genes in CP-A cells stably transfected with empty or ERBB2 expressing vectors grown in serum free media for 48 hrs. Non-specific IgG is provided as a control (n=3; ns= non-significant increases).

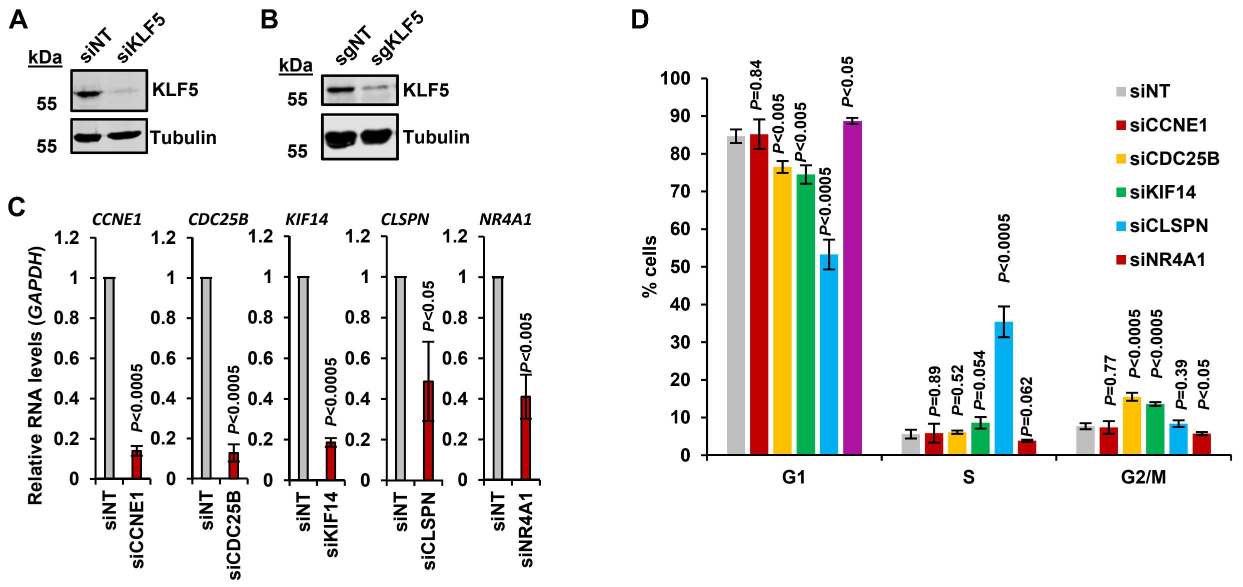


**Figure 5-figure supplement 3. KLF5 drives cell cycle progression in OE19 cells.** (A) Immunoblot of OE19 cell lysate from cells treated with either siNT or siKLF5 for 6 days and probed with antibodies against KLF5 and Tubulin. (B) Immunoblot of OE19-dCas9-KRAB lysate of OE19-dCas9-KRAB cells treated with a non-targeting guide or a pool of 3 guides targeting the KLF5 TSS and probed with antibodies against KLF5 and Tubulin. (C) RT-qPCR of RNA from OE19 cells treated with either siNT, siCCNE1, siCDC25B, siKIF14, siCLSPN or siNR4A1. The gene that was knocked-down was tested relative to *GAPDH* and *P*-values are shown (n=3). (D) Bar chart of the percentages of OE19 cells treated with either siNT, siCCNE1, siCDC25B, siKIF14, siCLSPN or siNR4A1 for 6 days in G1, S and G2/M phase of the cell cycle. *P-values* were calculated comparing each knockdown to siNT and *P-values* are shown (n=3).
