## Supplementary material for "Repurposing of KLF5 activates a cell cycle signature during the progression from a precursor state to Oesophageal Adenocarcinoma": Key Resource Table

**Template:**

| **Key Resources Table** | | | | |
| --- | --- | --- | --- | --- |
| **Reagent type (species) or resource** | **Designation** | **Source or reference** | **Identifiers** | **Additional information** |
| Cell line (*H. sapien*s) | OE19 | ACACC | 96071721 |  |
| Cell line (*H. sapiens*) | CP-A | ATCC | KR-42421 |  |
| Cell line (*H. sapiens*) | OE19-dCas9-KRAB | This study |  | OE19 transfected with vector to express dCas9-KRAB under doxycycline control |
| Cell line *(H. sapiens*) | CP-A-ERBB2 | This study |  | CP-A stably overexpressing ERBB2 |
| Cell line *(H. sapiens*) | CP-A-empty | This study |  | CP-A cells containing an empty vector control |
| Biological sample (*H. sapiens*) | Barrett’s oesophagus biopsies | Salford NHS FT |  | Freshly isolated from patients undergoing endoscopy |
| Biological sample (*H. sapiens*) | Oesophageal adenocarcinoma biopsies | Salford NHS FT |  | Freshly isolated from patients undergoing endoscopy |
| Transfected construct (human) | SmartPool siRNA against KLF5 | Horizon discovery | L-013571-00-0005 |  |
| Transfected construct (human) | SmartPool siRNA against ERBB2 | Horizon discovery | L-003126-00-0005 |  |
| Transfected construct (human) | SmartPool non-targeting siRNA | Horizon discovery | D-001810-10-0020 |  |
| Transfected construct (human) | Full length non-targeting guide RNA | Synthego |  | 5’-GUAAGGCUAUGAAGAGAUAC-3’ |
| Transfected construct (human) | Full length guide RNAs targeting KLF5 TSS | Synthego |  | 5’-GUGCGCUCGCGGUUCUCUCG-3’  5’-AGGACGUUGGCGUUUACGUG-3’  5’-GCGUCAAGUGUCAGUAGUCG-3’ |
| antibody | Rabbit monoclonal KLF5 antibody | Abcam | ab137676 | (1:10000) for western blot; 5ug for ChIP-seq |
| antibody | Mouse monoclonal tubulin antibody | Sigma-aldrich | T9026 | (1:2000) for western blot |
| antibody | Spike-in antibody | Active Motif | 61686 | 1ug for ChIP-seq |
| Antibody | Mouse monoclonal ErbB2 antibody | ThermoFisher | MA5-14057 | (1:1000) |
| Antibody | Mouse monoclonal AKT antibody | Cell signalling technology | 2920 | (1:2000) |
| Antibody | Rabbit monoclonal phosphor-Akt (S473) antibody | Cell signalling technology | 4060S | (1:2000) |
| Antibody | Rabbit monoclonal Erk1/2 antibody | Cell signalling technology | 4695S | (1:1000) |
| Antibody | Mouse monoclonal phosphor-Erk1/2 (T202,Y204) | Cell signalling technology | 9106S | (1:2000) |
| Antibody | Donkey anti-mouse secondary antibody (800CW) | Licor | 925-32212 | (1:10,000) |
| Antibody | Donkey anti-rabbit secondary antibody (700CW) | Licor | 925-32213 | (1:10,000) |
| Recombinant DNA reagent | pX330-U6-Chimeric_BB-CBh-hSpCas9 (plasmid) | Addgene | #42230 | AAVS guide RNA sequence 5’-GGGCCACTAGGGACAGGAT-3’ |
| Recombinant DNA reagent | pAAVS1-Puro-TRE-dCas9-KRAB-DNR (plasmid) | This study |  | pAS-4939 |
| Recombinant DNA reagent | pHAGE-ERBB2 | Addgene | 116734 |  |
| Recombinant DNA reagent | pHAGE-empty | This study |  | pAS-4940 |
| Recombinant DNA reagent | pMD2.G | Addgene | 12259 |  |
| Recombinant DNA reagent | psPAX2 | Addgene | 12260 |  |
| sequenced-based reagent | Primers | This study |  | See supplementary table S11 |
| commercial assay or kit | Lipofectamine™  RNAiMAX | Thermofisher | 13778150 |  |
| commercial assay or kit | Fugene HD | Promega | E2311 |  |
| commercial assay or kit | QuantiTect SYBR® Green RT-PCR Kit | Qiagen | 204243 |  |
| commercial assay or kit | RNeasy Plus Mini Kit | Qiagen | 74134 |  |
| commercial assay or kit | RNase-free DNase set | Qiagen | 79254 |  |
| commercial assay or kit | Ampure XP beads | Beckman Coulter Agencourt | A63881 |  |
| commercial assay or kit | TruSeq stranded RNA library kit v2 | Illumina | RS-122-2001 |  |
| commercial assay or kit | Nextera DNA library prep kit | Illumina | FC-121-1031 |  |
| commercial assay or kit | Nextera Index kit | Illumina | FC-121-1012 |  |
| commercial assay or kit | NEBNext high fidelity 2x PCR master mix | NEB | M0541 |  |
| commercial assay or kit | DNA Clean and Concentrator | Zymo | D4013 |  |
| commercial assay or kit | Polyfect | Qiagen | 301107 |  |
| commercial assay or kit | PEG-it | System Biosciences | LV810A-1 |  |
| commercial assay or kit | Polybrene | EMD Millipore | TR-1003 |  |
| chemical compound, drug | RS-1 | Sigma-Aldrich | R9782 | Used at final concentration 7.5 μM |
| chemical compound, drug | SCR7 pyrazine | Sigma-Aldrich | SML1546 | Used at final concentration 1 μM |
| chemical compound, drug | Doxycycline | Sigma-aldrich | D3447 | Used at final concentration of 100ng/mL |
| chemical compound, drug | propidium iodide | Sigma | P4170 | Used at 50 μg/mL |
| peptide, recombinant protein | RNase | Sigma | R4642 | Used at 100 μg/mL |
| peptide, recombinant protein | EGF | ThermoFisher | 10450-013 | 5 μg/L |
| peptide, recombinant protein | Bovine pituitary extract | ThermoFisher | 13028014 | Used at 50 mg/L |
| software, algorithm | Trimmomatic | Bolger et al, 2014 | V0.34 | http://www.usadellab.org/cms/?page=trimmomatic |
| software, algorithm | Bowtie2 | Langmean and Salzberg, 2012 | v2.3.0 | http://bowtie-bio.sourceforge.net/bowtie2/index.shtml |
| software, algorithm | Star | Dobin et al, 2013 | V2.5.4 | https://github.com/alexdobin/STAR |
| software, algorithm | Macs2 | Zhang et al, 2008 | v2.1.1 | https://github.com/taoliu/MACS |
| software, algorithm | Cufflinks | Trapnell et al, 2013 | v2.2.1 | http://cole-trapnell-lab.github.io/cufflinks/ |
| software, algorithm | DEseq2 | Love et al, 2014 | V1.22.2 | https://bioconductor.org/packages/release/bioc/html/DESeq2.html |
| software, algorithm | TOBIAS | Bentsen et al, 2020 | v0.5.1 | https://github.com/loosolab/TOBIAS |
| software, algorithm | featureCounts | Liao et al, 2014 | V1.6.2 | http://subread.sourceforge.net |
| software, algorithm | FastQC |  | v0.11.4 | https://www.bioinformatics.babraham.ac.uk/projects/fastqc/ |
| software, algorithm | bedtools | Quinlan and Hall, 2010 | v2.26.0 | https://bedtools.readthedocs.io/en/latest/ |
| software, algorithm | DeepTools | Ramírez et al, 2016 | V2.5.0 | https://deeptools.readthedocs.io/en/develop/ |
| software, algorithm | GSEA | Subramanian et al, 2005 | V3.0 | http://software.broadinstitute.org/cancer/software/gsea/wiki/index.php/Main_Page |
| software, algorithm | Homer | Heinz et al, 2010 | v4.9 | http://homer.ucsd.edu/homer/ |
| software, algorithm | R | R Core Team (2018) | v3.5.1 | https://www.r-project.org/ |
| software, algorithm | GraphPad Prism |  | V8.0 | www.graphpad.com |
| other | Crystal violet | Sigma Aldrich | HT90132 | Used at concentration of 0.1% |
| other | Gibco™ RPMI 1640 | ThermoFisher | 52400 |  |
| other | Gibco™ fetal bovine serum | ThermoFisher | 10270 |  |
| other | Gibco™ penicillin/streptomycin | ThermoFisher | 15140122 |  |
| other | Keratinocyte SFM (1x) | ThermoFisher | 17005042 |  |
